# Evidence of Activity-Silent Working Memory in Prefrontal Cortex

**DOI:** 10.1101/2024.06.03.597259

**Authors:** Weicheng Tao, Camilo Libedinsky

**Affiliations:** National University of Singapore

## Abstract

The lateral prefrontal cortex encodes working memory and motor preparation information, but the underlying neural mechanisms are debated. Recurrent neural network models relying on persistent neural activity have been challenged by the observation of periods of absent activity and information during memory maintenance, implying the existence of activity-silent mechanisms. To assess whether activity-silent mechanisms are needed for working memory maintenance, we recorded neural activity in macaque prefrontal cortex during a delayed-saccade task. We replicated the observation of periods of absent activity and decreased information between bursts of gamma power, but we show that these results are consistent with models that rely exclusively on persistent activity. However, an assessment of the length of periods with absent selective activity across the population revealed that activity-silent mechanisms are indeed needed to maintain memory information, although this is only evidenced in a small fraction of trials.

## Introduction

The ability to store and manipulate information about past stimuli and upcoming movements (working memory and motor preparation, respectively) is fundamental for intelligent behavior. Regions of the dorsolateral prefrontal cortex are involved in both of these cognitive operations (Funahashi et al., 1989; Fuster & Alexander, 1971; Jonikaitis et al., 2023), and some prefrontal regions can encode both types of information simultaneously (Tang et al., 2020).

Recurrent neural network models with attractor states have been postulated as a mechanism to maintain information in the absence of inputs (Compte et al., 2000; X.-J. Wang, 2021). In these models, neurons with similar selectivity excite each other recurrently (either directly or indirectly) such that activity across the population of selective neurons persists over time without external inputs to the network. This recurrent excitation can generate “attractor states”: when the network undergoes minor deviations from one of these states (e.g., due to noise or weak inputs), it evolves back towards the original state over time. Physiological evidence suggests that prefrontal regions maintain information in attractor states (Inagaki et al., 2019; Parthasarathy et al., 2019; S. Wang et al., 2023; Wimmer et al., 2014).

Recurrent neural networks that rely purely on the activity of neurons to maintain their attractor states (activity-RNNs) require one or more “active” neurons at every point during the memory maintenance period; otherwise, the network would lose the memory. For a neuron to be active, it needs to fire an action potential, and the effect of an action potential on postsynaptic neurons lasts for several milliseconds due to the temporal filters imposed by synaptic transmission (Szűcs, 1998). For example, the slow dynamics of NMDA receptors (∼100 ms decay) are essential to sustain activity in attractor network models (Compte et al., 2000, p. 200; X.-J. Wang, 1999). Thus, activity-RNNs lose their information if the neurons in the network are silent for longer than ∼100 ms.

The ability of activity-RNNs to model dorsolateral prefrontal activity has been questioned based on the observation that during the retention of working memory information, prefrontal neurons have synchronized periods of silence, during which the overall activity of the population decreases to baseline levels and memory information cannot be decoded (Barbosa et al., 2020; Lewis-Peacock et al., 2012; Lundqvist et al., 2016; Miller et al., 2018; Panichello et al., 2023; Rose et al., 2016; Shafi et al., 2007; Sprague et al., 2016; Stokes, 2015; Wolff et al., 2017). Thus, activity-RNNs were updated by adding short-term plasticity (STP) to their synapses (stp-RNNs) (Masse et al., 2019; Mikael Lundqvist, 2011; Mongillo et al., 2008). STP is the transient change of synaptic efficacy over tens of milliseconds to minutes, and it is known to exist in the prefrontal cortex under certain conditions (Fujisawa et al., 2008; Hempel et al., 2000). These stp-RNN models maintain information using both activity and STP selectively during the activity-silent periods.

While activity-silent memory periods are the primary motivation behind stp-RNN models, their existence is debatable. For instance, activity-silent memory periods could result from measuring the activity in brain regions where the information is not maintained (Constantinidis et al., 2018). It could also result from measuring brain activity using techniques that lack the spatial resolution needed to identify the selective neurons that show elevated activity (Constantinidis et al., 2018). Despite these criticisms, two lines of evidence suggest that activity-silent memory periods exist: (1) informative spiking is exclusively present in gamma burst periods (Lundqvist et al., 2016), and (2) decoding of memory information in single trials can drop to chance levels for extended periods (>100 ms), even when recording from a large number of neurons (>300) (Panichello et al., 2023). Both observations rely on *decreased* activity for extended periods such that information is lost. However, a decrease in activity or information is insufficient to lose memory information in activity-RNNs since even a low number of spikes can sustain a memory. If the population of *selectively* active neurons was silent for extended periods (> ∼100 ms), that would be a strong argument against activity-RNNs, and in favor of stp-RNNs. Such coordinated periods of silence have been observed in the cortex in other contexts (i.e., OFF periods during slow-wave sleep) (Vyazovskiy et al., 2011), so they may also exist during memory maintenance in the prefrontal cortex.

Lundqvist and colleagues (2016) found decreased activity and information during off-gamma periods, and they interpreted these observations as supporting the existence of activity-silent memory maintenance mechanisms during these off-gamma periods. However, for this evidence to support activity-silent memory maintenance mechanisms, information during off-gamma periods should be absent (not just reduced). Thus, we hypothesized that there would be no memory information during off-gamma periods. Further, since Panichello and colleagues (2023) found chance level decoding of working memory information in single trials for extended periods (> 100 ms), we hypothesized that there would be time periods during which selective spiking is absent (i.e., activity-silent periods). Our results did not support the first hypothesis, but we did find evidence in support of the second. An analysis of the length of activity-silent periods across selective neurons revealed that some periods were longer than expected by chance, implying that activity-silent mechanisms must be at play. Hence, we conclude that STP is necessary to explain the maintenance of working memory and/or motor preparation in the prefrontal cortex.

## Methods

### Subjects and recordings

We recorded brain activity in two male adult macaques (*Macaca fascicularis*, Subject P and J) while they performed Task A and a third one (Subject W) while he performed Task B (Figure 1). All animal procedures were approved by, and conducted in compliance with, the standard of the Agri-Food and Veterinary Authority of Singapore and Singapore Health Services Institutional Animal Care and Use Committee (SingHealth IACUC #2012/SHS/757), and the National University of Singapore Institutional Animal Care and Use Committee (NUS IACUC #R18-0295). Additionally, these procedures followed the recommendations outlined in the *Guidelines for the Care and Use of Mammals in Neuroscience and Behavioral Research* (National Academies Press, 2003).

**Figure 1.**
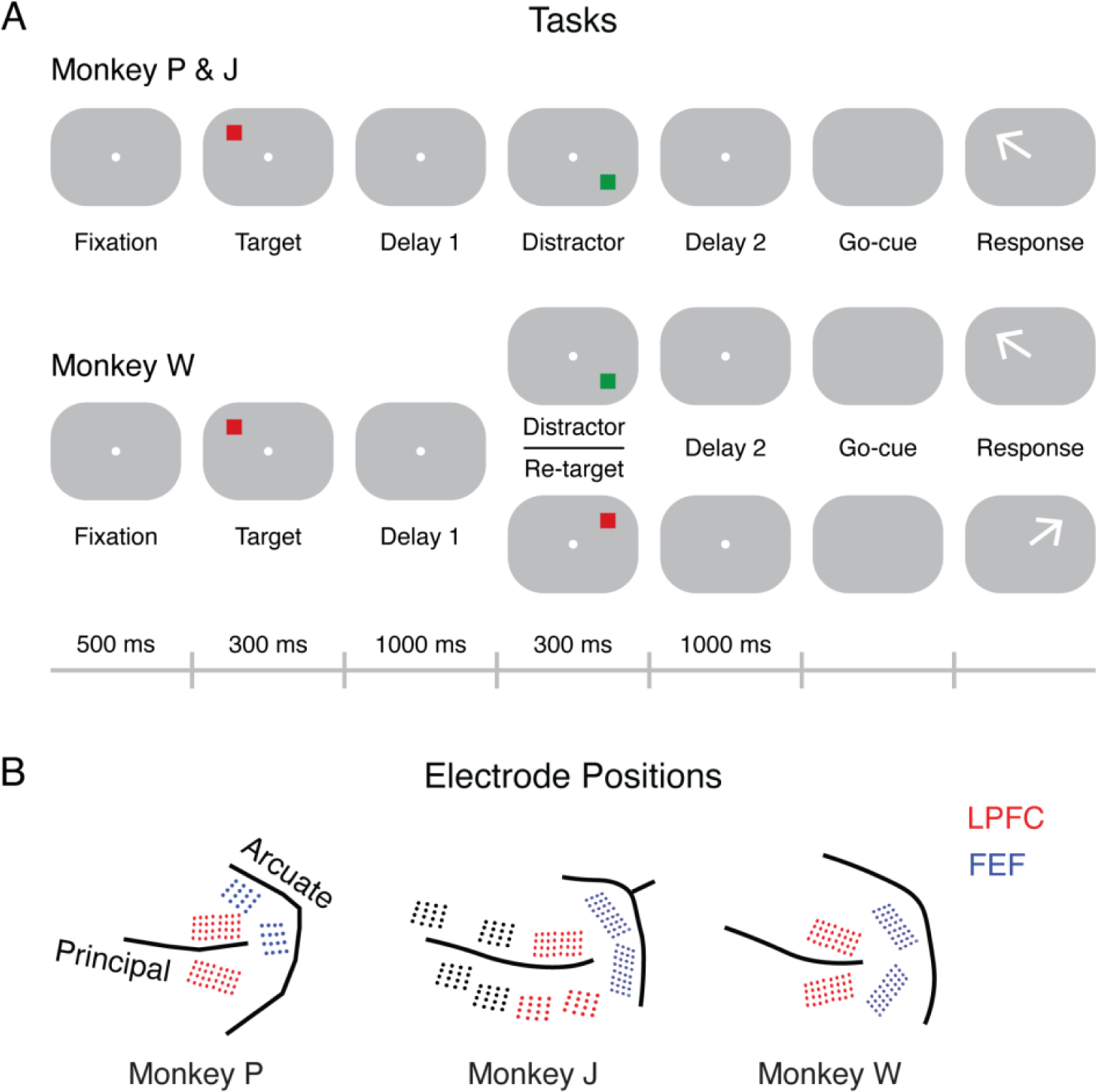
Delayed-saccade spatial WM task with intervening distractor or new target. **(A)** Tasks performed by monkeys P and J (top), and monkey W (bottom). Task A and B have similar trial structures except that the intervening stimulus is always a distractor in Task A, while it can be a new target in Task B in half of the trials. **(B)** Electrode locations for the three monkeys (FEF: blue; LPFC: red; not included in the analysis: black).

We implanted a titanium head post (Crist Instruments, MD, USA) and intracortical microelectrode arrays (Microprobes, MD, USA) only in the left hemisphere of each subject. Subject P had two 32-electrode arrays in Area 9/46 (LPFC) and two 16-electrode arrays in Area 8A (FEF). Subject J had four 16-electrode arrays in Area 46 (not included in the analysis), one 32-electrode array and two 16-electrode arrays in Area 9/46 (LPFC) and two 32-electrode arrays in Area 8A (FEF). Subject W had two 32-electrode arrays in Area 9/46 (LPFC) and two 32-electrode arrays in Area 8A (FEF). The arrays were made of platinum-iridium wires, separated by either 200 or 400 μm, ranging from 1 to 5.5 mm in length (we inserted longer electrodes into the sulci), and had an impedance of 0.5 MΩ. These wires are arranged in 4 by 4 or 8 by 4 grids.

We recorded neural signals in subjects P and J using a 128 or 256-channel Plexon OmniPlex system (Plexon Inc., TX, USA) at 40 kHz sampling rate. The neural signals in subject W were recorded using a 128-channel Grapevine recording system (Ripple Neuro, UT, USA) at 30 kHz sampling rate. We recorded the eye positions of each subject using the Eyelink 1000 Plus (SR Research Ltd, ON, CA). The same computer was used to simultaneously record the neural signals and the eye positions. We designed the behavioral tasks using the Psychtoolbox in MATLAB (Mathworks, MA, USA) or Psychopy in Python. During the recording, we ran the tasks on a standalone computer connected to the recording computer using parallel ports for event mark synchronization.

### Behavioral tasks

For Monkeys P and J, we presented targets and distractors on a 3 x 3 grid (at 10 degrees of visual angle). After a 500 ms pre-target fixation period, we flashed a red square (the target) for 300 ms in one of the eight outer positions on the grid. A 1s delay period followed (Delay 1), after which we flashed a green square (the distractor) for 300ms in one of 7 positions (target and distractor never occupied the same position). Another 1s delay period followed (Delay 2) before the disappearance of the central fixation spot, which indicated to the animals that they should respond with an eye movement to the remembered target position. To receive the reward, constituting a drop of fruit juice, the animal had to maintain fixation on the target position for 200 ms (Fig. 1A top).

The trial structure for Monkey W was similar but had three key differences. First, the animals were presented with crosses at the target positions instead of a grid outline indicating the possible target positions. Second, we only presented the animals with 4 possible target positions, rather than the 8 presented to Monkeys P and J. Third, the second stimulus could either be a new target (50% chance), or a distractor (50% chance), rather than only distractors (Fig. 1A bottom).

We only used the corner positions when comparing monkeys J’s and P’s results with monkey W’s. In other words, we did not include data from targets located vertically and horizontally for monkeys P and J. Monkey J did not have enough trials for the bottom-right target location, so we used the target in the middle-right instead.

### Signal processing

All initial neural recordings were bandpass-filtered between 300 and 3,000 Hz for spike sorting. The single-unit activities were extracted from each channel separately using an automated spike sorting algorithm based on the Hidden Markov model. We extracted spikes from each trial and computed the firing rates using a 50 ms moving window (5 ms per step) for each identified single unit. We also computed the firing rate of a baseline period (-400 to 0 ms relative to the target onset) for each trial. We then computed the normalized firing rates using the baseline firing rates’ mean and standard deviation (SD) across trials.

The initial recordings were also bandpass-filtered between 10 and 150 Hz and downsampled to 1,000 Hz to generate LFPs for analyses of oscillatory activities. We used a multi-taper method to estimate the power spectrogram based on single-trial LFPs. In particular, we used the discrete prolate spheroidal sequences (DPSS) to generate tapered window functions that are orthogonal with each other (Percival & Walden, 1993). These window functions were convolved with the LFPs separately, and the results were transformed into power spectrograms using the short-time Fourier transform (STFT) with 1 ms temporal resolution and 1 Hz frequency resolution. The resulting power spectrograms were then normalized to power spectral density (PSD) and averaged across window functions to acquire a single estimate per trial. We set the window width (N) to 100 ms and the half-bandwidth (W) to 15 Hz. The DPSS would generate two window functions with these settings to provide the optimized spectral estimation.

The 1/f normalization was applied to the spectrograms to compensate for the 1/f spectral structure of brain dynamics. The resulting spectrogram of each trial was dB converted per frequency relative to the mean power of a baseline period (-400 to 0 ms relative to the target onset). These dB-converted spectrograms were then averaged across trials and channels to generate the mean spectrogram. The spectrograms were also averaged across frequencies in the gamma band (40 to 120 Hz) to generate time-varying gamma power.

### Percentage of explained variance

To quantify the target discrimination exhibited in neural activities, we estimated the percentage of explained variance (PEV) in terms of ω^2^. The ω^2^ was computed based on neural activities recorded for targets in different locations (i.e., groups). It measures how much of the variance in neural activities can be explained by the distinction across groups. The SS, MS, and df are the sum of squares, mean square, and degree of freedom, respectively.

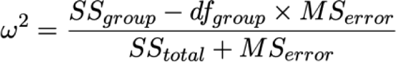

We used the time-varying firing rates of each unit to calculate the time-varying PEV. They were then averaged across units to calculate the mean PEV. Note that the ω^2^ can sometimes be a small negative number. If the ω^2^ is negative, then the target location cannot explain the activity variance.

### Selective units and channels

To identify selective units, we compared the time-varying PEV of each unit with a predefined PEV threshold (5% explained variance). If a unit exhibited high PEV (higher than the threshold) in two adjacent 50 ms time bins before the distractor onset (between 0 ms and 1300 ms after target onset), the unit was considered a selective unit. We defined a selective channel as any channel from which we recorded at least one selective unit. Unless specified otherwise, we only analyzed selective units and channels in the following analyses.

### Response rank and preferred target location

For each selective unit, we estimated the mean firing rate during the target presentation period (100 to 300 ms relative to the target onset) for each target location. Then, we ranked the target locations based on their mean firing rates. R1 was used to label the location with the highest firing rate (i.e., the preferred target location), and R4 was used to label the location with the lowest firing rate.

### Decoding performance

In addition to the PEV, we also estimated the maintained target information using a support vector machine (SVM) on individual neurons (note that this is single-neuron decoding and not population decoding). We trained SVM classifiers for each selective unit and timepoint using the firing rates and the corresponding target locations and estimated the decoding performance in terms of the f1-score. More specifically, 1) the firing rates of all trials were randomly split into two partitions, one with 60% of trials (train set) and the other one with 40% of trials (test set); 2) both sets were normalized using the mean and SD of the train set; 3) an SVM classifier (linear kernel with the regularization parameter C = 1) was trained on the train set and tested on the test set; 4) the test results were used to compute the true positive (TP), false positive (FP) and false negative (FN) for each target location; 5) The TP, FP and FN were used to compute the f1-score for each target; 6) the resulting f1-scores were averaged across targets; and finally, 7) repeating steps 1-6 fifty times and averaging the results.

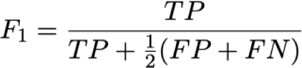

### Population inter-spike interval

In each brain area (LPFC or FEF), simultaneously recorded selective units were used to estimate the duration of coordinated activity-silent periods across populations with overlapping selectivity. We combined spikes extracted from two or more selective units sharing the same preferred target location. While tens of units could be recorded simultaneously in a recording session, only a fraction of them would show overlapping selectivity. We could generate at most four sets of combined activities (one for each target location) for each recording session. We then computed the population inter-spike intervals (population-ISIs) for each combined activity. We analyzed a total of 16 sessions, which means that there were a maximum of 128 possible empirical distributions (2 areas x 4 locations x 16 sessions). However, since not all sessions/locations had multiple neurons with overlapping selectivities, we analyzed the 39 sessions/locations that fulfilled that criterion. Further analyses were conducted based on these 39 empirical distributions. For each trial, we analyzed the activity during the Delay 1 period (500 - 1200 ms after the go-cue).

### Null distribution of the maximum population-ISI

To determine whether activity-silent periods in the data were longer than expected by chance we generated a distribution of random population-ISIs by shuffling trials. We extracted one maximum population-ISI from 2000 shuffles to generate a null distribution of the maximum population-ISIs (null-max-ISI). This distribution was then compared to the empirical-max-ISI to determine whether it was expected by chance. Empirical-max-ISIs that exceeded the 99^th^ percentile were considered significantly larger than chance.

### Gamma modulation

Selective activities were suggested to exclusively appear at gamma modulated (gamma-mod) channels (Lundqvist et al., 2016). To identify gamma-mod channels, we compared the gamma power during the target period (100 – 300 ms) with a baseline period (-200 – 0 ms). If target period showed significantly higher gamma power (Wilcoxon signed-rank test), the channel was considered a gamma-mod channel. In addition, the units extracted from gamma-mod channels were considered gamma-mod units. We included both selective and non-selective units in gamma-mod related analyses to test whether WM related selective activities appeared exclusively at gamma-mod channels.

### Gamma burst and off-gamma periods

Enhanced gamma modulation could be manifested as increased number of intermittent gamma bursts (Lundqvist et al., 2016). As existing studies suggested, the gamma bursts may occur at narrow frequency bands instead of spanning the entire gamma band (Lundqvist et al., 2018). These studies performed the burst extraction procedure on pre-defined gamma sub-bands (Lundqvist et al., 2016; Lundqvist et al., 2018). However, since this procedure is sensitive to how the sub-bands are defined, we modified the burst extractions as follows: we performed the burst extraction procedure for each gamma band frequency (40, 41 … 120 Hz) separately and then aggregated the results across frequencies. More specifically, for each trial and gamma band frequency, we defined a power threshold and a duration threshold. We used the 90^th^ percentile of the power as the power threshold and three cycles of that frequency to be the duration threshold. These configurations allowed us to identify high gamma power periods and reduce the false positive rate. Across gamma band frequencies, a period was considered a gamma burst if its power passed both power and duration thresholds in any of the gamma frequencies.

In order to evaluate the effect of gamma bursts on neuronal selective activities, we also identified off-gamma periods in each trial. A period was considered an off-gamma period if its power did not pass the power threshold and lasted longer than 25 ms (equivalent to three cycles of 120 Hz oscillation) in all the gamma frequencies. The gamma burst and off-gamma period could never overlap, and there were some periods that were neither gamma burst nor off-gamma periods.

In individual trials, we binary-coded burst period time points to be ones and the rest of the time points to be zeros. They were then averaged across trials to generate the time-varying gamma burst rate. A similar approach was used to generate the rate of off-gamma period.

### Statistical methods

The time-varying metrics included firing rate, PEV, decoding performance, gamma power, gamma burst and off-gamma rates. These time-varying metrics were plotted with the standard error (SE) estimated using bootstrapping. When comparing the same measurement across two conditions, such as LPFC vs. FEF or Delay 1 vs. Baseline, we used permutation to test the mean difference between conditions. The permutation test was also used to compare a measurement against 0 (assigning random signs to the data to generate a null distribution of the metric).

To determine whether the decoding performance was higher than chance level, the location labels were shuffled before model training to estimate the chance level decoding performance (averaged across 50 shuffles). It was then compared with the empirical decoding performance using permutation test.

To compare the empirical population-ISIs with the null-max-ISI distribution, we checked whether there were empirical ISIs that were longer than the 99^th^ percentile (duration threshold) of the null-max-ISI distribution. To account for the family-wise error rate, Bonferroni correction was used to adjust the threshold.

## Results

### Delay period activity and single neuron information

We recorded the activity of 428 single units across 16 sessions in 3 monkeys trained to perform a delayed saccade task with an interfering distractor. While the FEF contained a higher proportion of selective neurons during the cue and first delay period (48% vs. 26% of neurons, Table 1), selective neurons in the LPFC were, on average, more informative during the Delay 1 period (Figure 2A, p = .010, permutation test; 500 to 1200 ms after target onset). We corroborated this observation using single-neuron decoding analyses (Figure 2B, *p* < .001).

**Table 1.** The number of selective and non-selective single units (and the number of channels from which these units were recorded) in LPFC (top) and FEF (bottom). Selectivity was measured during the Cue and Delay 1 period. A breakdown per monkey can be found in Table S1.

| Area | Selectivity | Units | Channels |
| --- | --- | --- | --- |
| LPFC | Selective | 50 (26%) | 37 (27%) |
|  | Non-selective | 140 (74%) | 101 (73%) |
| FEF | Selective | 114 (48%) | 88 (52%) |
|  | Non-selective | 124 (52%) | 80 (48%) |

**Figure 2.**
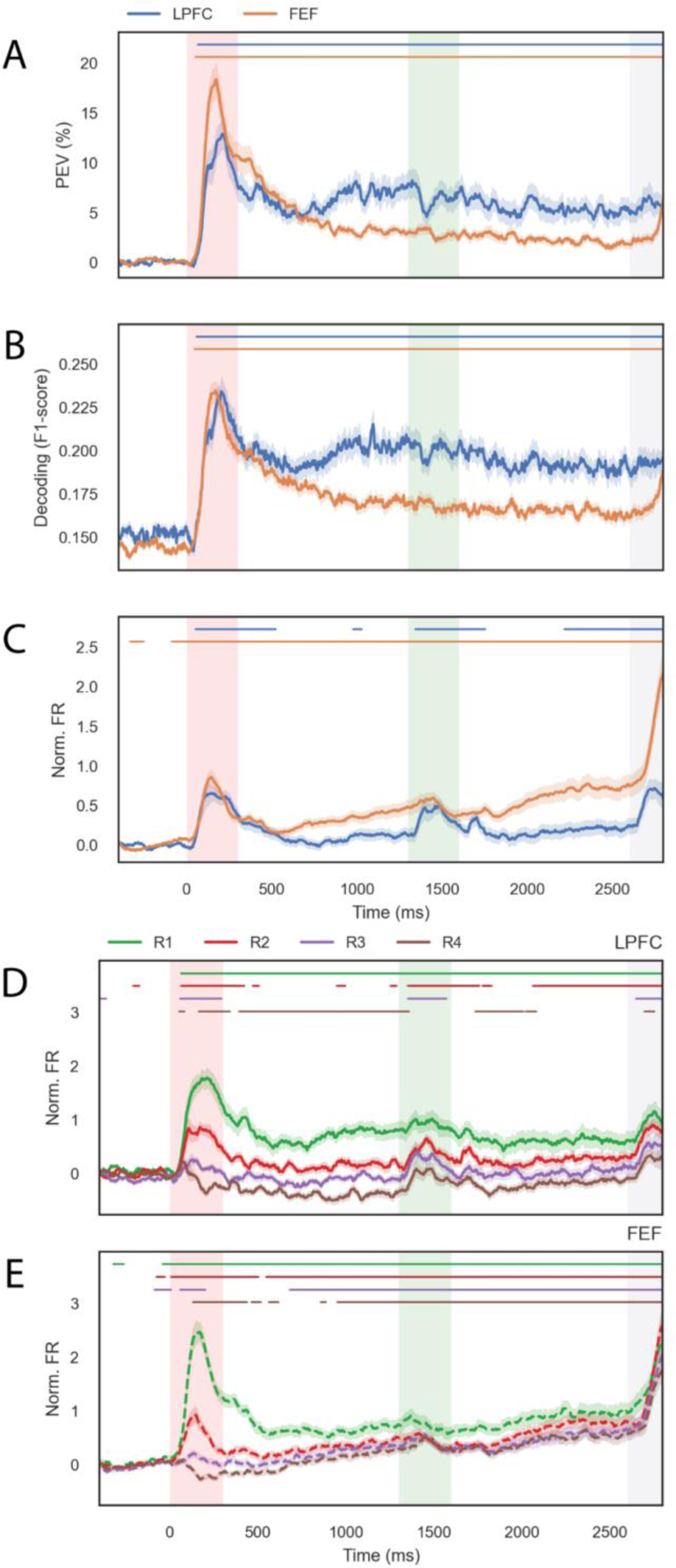
Single-neuron activity and information. **(A)** Average percentage of explained variance (PEV) across selective neurons for LPFC (blue) and FEF (orange). Shaded areas correspond to the SEM (estimated using bootstrapping). **(B)** and **(C)**, the same convention as in (A), but for the average decoding performance (F1-score) of single neurons and the average normalized firing rate, respectively. **(D)** Average normalized firing rate of LPFC neurons separated by activity rank (R1 - R4) during the target period (where R1 corresponds to the location with the highest average activity, and R4 corresponds to the location with the lowest average activity). **(E)** Same as (D), but for FEF neurons. For all plots, we highlight the target presentation period in red shade (0 m - 300 ms), the distractor presentation period in green shade (1300 ms - 1600 ms), and the response period in gray (go-cue at 2600 ms). For plot (B), we show significance against a label-shuffled F1-score. For the rest of the plots, we show significance against the baseline with the solid lines above the plots (uncorrected *p* < .05 for 5 consecutive bins).

Previous studies have shown that the LPFC has periods of baseline-level activity during memory maintenance, consistent with activity-silent mechanisms (Wolff et al., 2017). A recent study showed that averaging across all neurons masked neurons that generated persistent activity selective to their preferred stimuli, which carried most of the information (Thrower et al., 2023). We confirmed that the average activity of selective neurons in the LPFC dropped to baseline during the delay periods (Figure 2C, p = .073, permutation test). However, this was not the case in the FEF (Figure 2C, p < .001). To understand why the average activity in LPFC dropped to baseline levels, but the information was high, we replotted the mean normalized average neuron activity separately for different locations. For each neuron, we determined the rank order of locations, such that R1 corresponded to the location that elicited the highest average firing rate, R2 the second highest, R3 the third highest, and R4 the lowest. The average of R1 responses of LPFC neurons remained consistently above baseline during the Delay 1 period (Figure 2D green line, p < 0.001), while the average R4 responses remained below baseline during the same period (Figure 2D, brown line, p < 0.001). In contrast, the average activity of FEF neurons for all ranks was elevated above baseline (Figure 2E, p < 0.001) except R4 (p = .081) and ramped up over time (for single monkey analyses, see Table S1 and Figure S1).

These analyses show that baseline-level population activity during delay periods is consistent with selective neurons showing elevated activity. These results highlight the importance of considering the selectivity of neurons when averaging across them. Thus, we employ this approach in subsequent analyses.

### Activity and information during gamma-bursts and off-gamma periods

It has been proposed that gamma bursts flank periods of activity-silent memory maintenance (Lundqvist et al., 2016). Thus, we hypothesized that information would be absent during off-gamma periods. Before directly testing this hypothesis, we assessed whether our data could replicate the main results reported by Lundqvist and colleagues (2016). Indeed, we could replicate all of their key results. We confirmed that informative spiking is exclusively present in electrodes that contain gamma bursts (i.e., gamma-modulated electrodes)(Figure 3A-B). We also confirmed that gamma power and gamma-burst rate increased during stimulus presentation and remained elevated during the delay period (Figure 3C). We found that the off-burst rate decreased during the stimulus presentation period but remained at baseline levels during the delay period (Figure 3C). Finally, we confirmed that during gamma-burst periods, the activity of neurons was higher than during off-periods (Figure 3D).

**Figure 3.**
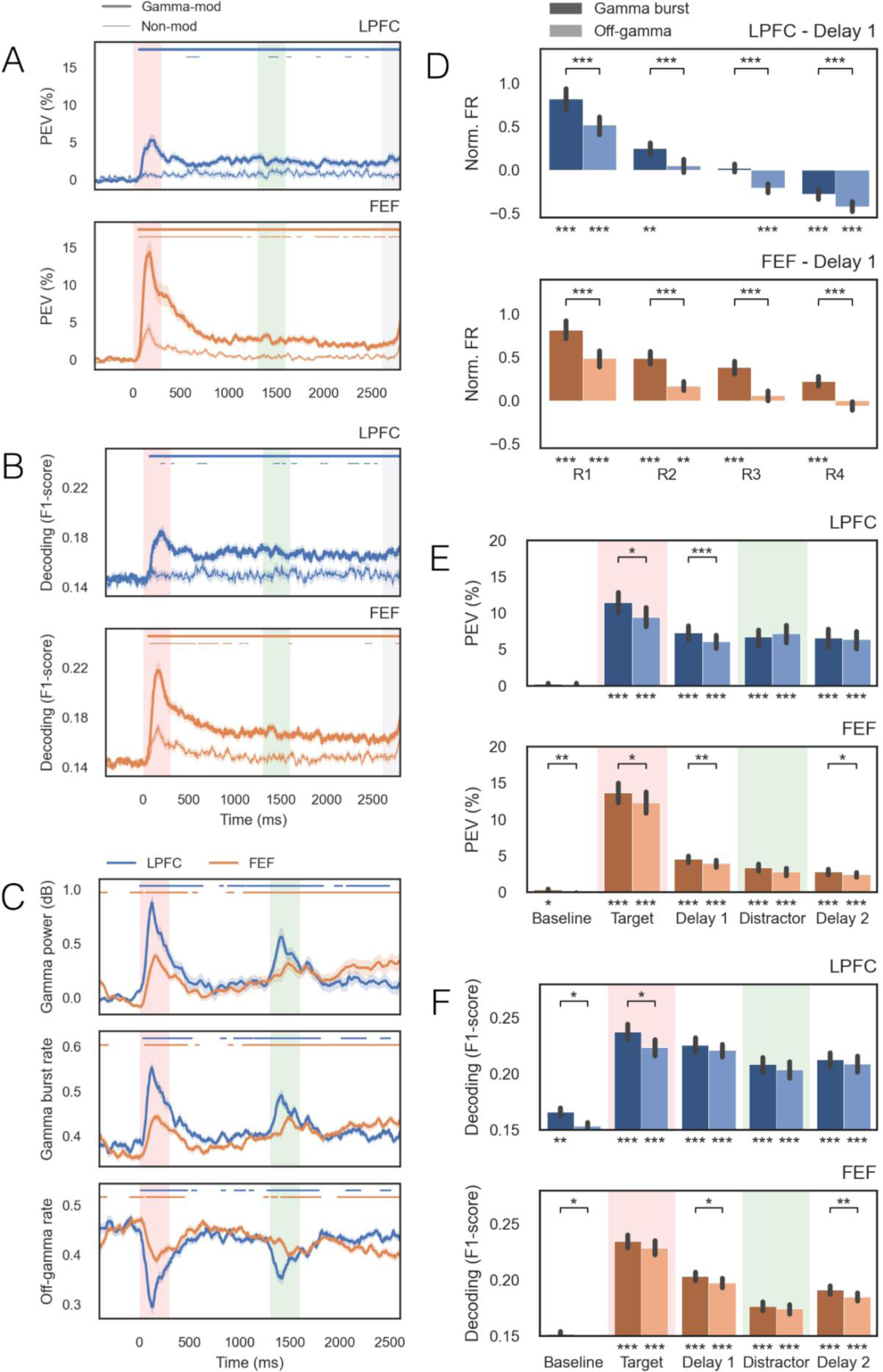
Activity and information inside and outside gamma bursts. ***(A)*** PEV as a function of time plotted separately for neurons recorded from gamma-modulated electrodes (thick lines) and non-modulated electrodes (thin lines) in the LPFC (blue) and FEF (orange). We included all neurons (selective and non-selective). **(B)** Average single-neuron decoding performance of target location. **(C)** Gamma power (top), gamma burst rate (middle), and off-gamma burst rate (bottom) as a function of time. **(D)** Normalized average firing rate of selective neurons during the Delay 1 period for gamma-burst (dark colors) and off-gamma burst (light colors) periods separated by selectivity rank. **(E)** Mean PEV of the activity within gamma-bust (dark colors), and off-gamma burst (light colors) periods for LPFC (blue) and FEF (orange). **(F)** Same convention as in (E) but for the performance of single neuron decoding of memory information. For all plots, we highlight the target presentation period in red (0 m - 300 ms) and the distractor presentation period in green (1300 ms - 1600 ms). For plot (B), we show significance against a label-shuffled F1-score. For (A) and (C), we show significance against the baseline with the lines above the plots (uncorrected p < 0.05 for 5 consecutive bins). For (D) to (F), we show the significance against the baseline with asterisks below the plots and the significance between gamma-burst and off-gamma periods with asterisks above the brackets (*** = p < 0.001, ** = p < 0.01. * = p < 0.05)

These results were taken as support for the view that “… WM information is only present in brief bursts of spiking and maintained in synaptic changes between such events” (Lundqvist et al., 2016). This conclusion rests on the assumption that working memory information is present in the neuronal spiking during gamma bursts but absent during off-gamma periods (during which memory maintenance would require STP). Thus, we hypothesized that there would be no memory information during off-gamma periods. To test this hypothesis, we calculated single-neuron information during the delay period within gamma-burst and off-gamma periods (Figure 3E-F). During the Delay 1 period, we found more information within gamma bursts than off-bursts (Figure 4E-F; p>0.05). However, during the Delay 1 period, the information during off-gamma periods was higher than chance (Figure 4E-F; p<0.001) (for single monkey analyses, see Table S1 and Figure S1). Thus, our results did not support our first hypothesis that working memory information is present in neuronal spiking during gamma bursts but absent during off-gamma periods. Hence, these results do not support the view that STP is necessary for working memory maintenance during off-burst periods.

**Figure 4.**
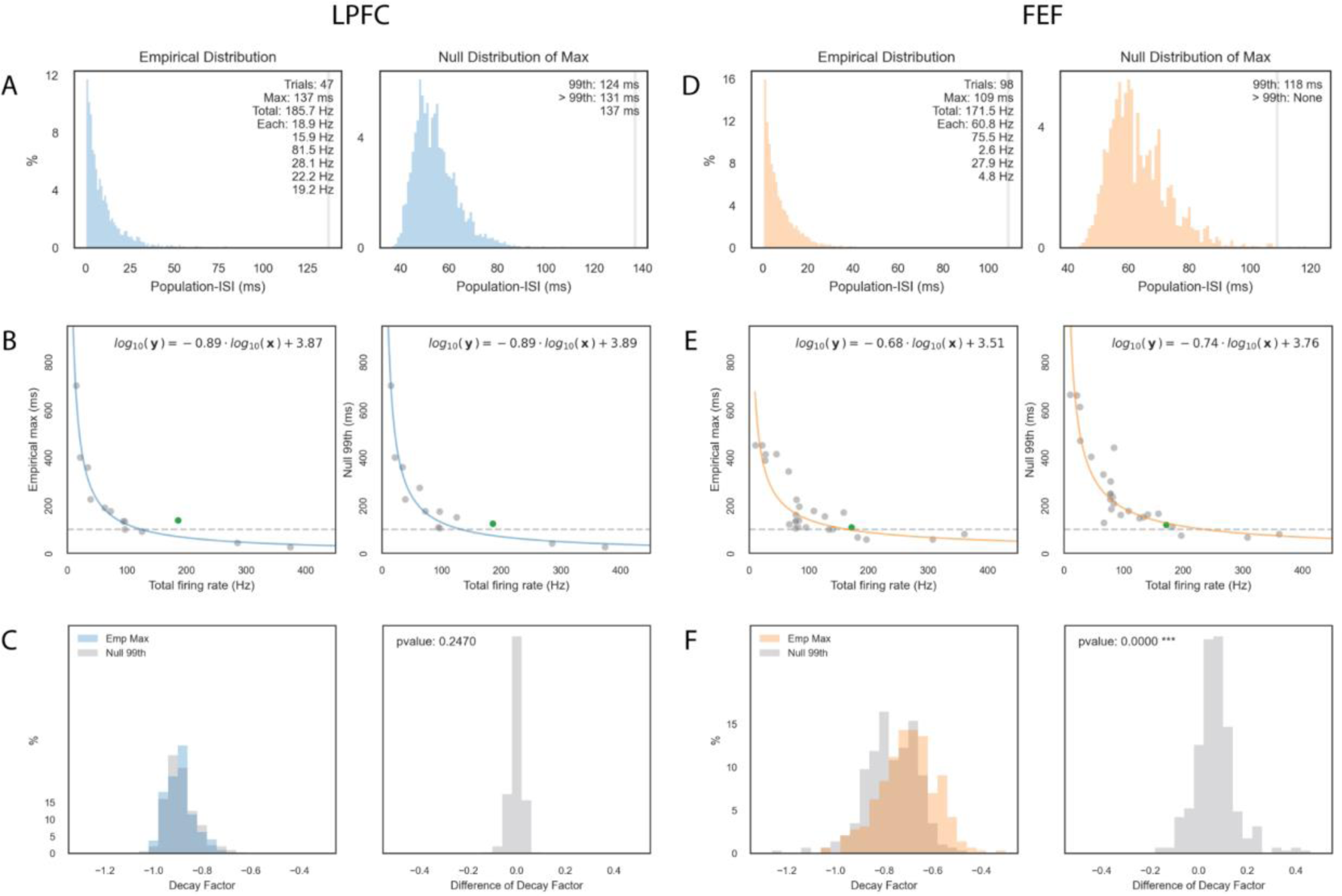
Length of activity-silent periods in single trials. **(A)** *(left)* An example of an empirical-max-ISI distribution for LPFC units with the same target selectivity recorded simultaneously in a single session. The gray vertical bar indicates the maximum empirical-max-ISI observed across all trials. *(right)* The distribution of the null-max-ISI was computed by shuffling the trials of individual units. The gray vertical bar indicates the 99th percentile of the null-max-ISI. **(B)** *(left)* The maximum empirical-max-ISI plotted against the total firing rate for each session and location. The line represents the exponential model fit, and the horizontal dotted line shows the 100 ms line. The green dot shows the example distribution shown in (A). *(right)* Same as (*left)* but for the null-max-ISI distributions. **(C)** *(left)* Distribution of decay factors calculated by bootstrapping the sessions/locations (selecting 10) for empirical-max-ISI fits (blue) and null-max-ISI fits (gray). *(right)* Distribution of the difference between decay factors of empirical minus null-max-ISIs (one difference for each bootstrap). **(D-F)** Same as (A-C), but for FEF.

### Population-level activity-silent periods

Panichello and colleagues (2023) reported long periods (> 100 ms) during the delay period where decoding of working memory information was at chance level (Panichello et al., 2023). Based on this report, we hypothesized that the chance level of decoding they observed was due to activity-silent periods, reflected in an absence of spikes across the population of selective neurons.

To test this hypothesis, we measured the duration of silent periods in single trials across all the simultaneously-recorded neurons with overlapping selectivity. First, we pooled the spike times of all the neurons with overlapping selectivity. Then, we calculated the interspike interval for every pair of adjacent spikes in this population (population-ISI). This population-ISI serves as an upper bound to the activity-silent periods in the whole population. Figure 4A shows an example distribution of population-ISI in a population of 6 LFPC neurons with overlapping selectivity recorded simultaneously and with a population firing rate (population-FR) of 185.7 Hz. For each distribution, we identified the largest population-ISI in the session (empirical-max-ISI)(Figure 4A left, 137 ms, vertical gray line). To determine whether the empirical-max-ISI was larger than the max-ISI expected by chance, we measured the max-ISI of data after shuffling trials (e.g., we pooled the activity of cell A in trial X with the activity of cell B in trial Y). We repeated this process 2000 times to generate a null distribution of max-ISIs (Figure 4A right). In both FEF and LPFC, the empirical-max-ISIs and the null-max-ISIs decreased exponentially as the population-FR increased (Figure 4B, E). If activity-silent periods exist, we would expect the empirical distribution exponential decay to be slower than that of the null distribution since long ISIs would exist in the empirical data but not in the shuffled data. Consistent with the existence of activity-silent periods, the exponential decay of empirical distribution in FEF was significantly slower than the null distribution (*p* < 0.001, Figure 4F). However, they were not different in the LPFC (*p* > 0.01, Figure 4C).

A more explicit test of the existence of activity-silent periods is to determine whether individual population-ISIs exceed the length of those expected by chance. Across LPFC and FEF, we observed 9 out of 39 empirical distributions (23%) in which at least 1 empirical-max-ISI exceeded the 99th percentile of the shuffled distribution (LPFC: 4 out of 13, 31%; FEF: 5 out of 26, 19%; Figure 5). For those 9 distributions, 10 silent periods (0.007%) exceeded the 99th percentile of the shuffle, ranging from 43 to 414 ms (9 out of these 10 silent periods were larger than 100 ms). Even though these events are rare, these results are consistent with the existence of activity-silent periods.

**Figure 5.**
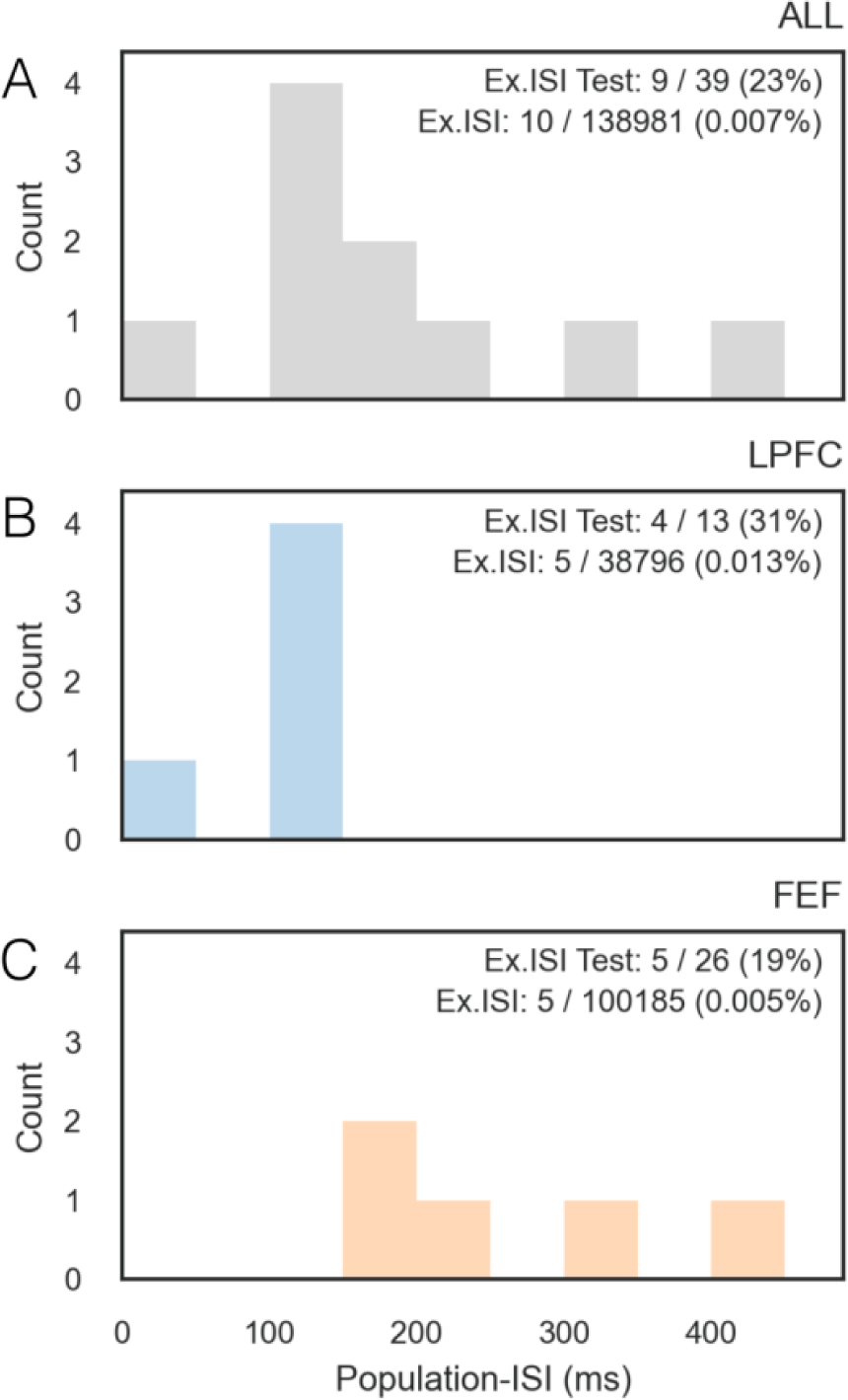
LPFC and FEF have longer activity-silent periods than expected by chance. **(A)** The number of activity-silent periods plotted against their length (in ms) for both regions combined, **(B)** for LPFC and **(C)** for FEF.

## Discussion

Recurrent neural network models that rely exclusively on the activity of neurons (activity-RNNs) can maintain memory information in the absence of stimuli. However, recent studies have shown that LPFC networks have coordinated periods of decreased spiking and absence of information (Barbosa et al., 2020; Lewis-Peacock et al., 2012; Lundqvist et al., 2016; Miller et al., 2018; Panichello et al., 2023; Rose et al., 2016; Shafi et al., 2007; Sprague et al., 2016; Stokes, 2015; Wolff et al., 2017). To explain memory maintenance without spiking, activity-RNN models were updated to include short-term synaptic plasticity (stp-RNNs) (Masse et al., 2019; Mikael Lundqvist, 2011; Mongillo et al., 2008). Here, we tested three key assumptions of stp-RNN models. The first assumption is that trial-averaged activity decreases to baseline during memory maintenance. The second assumption is that spiking information is uninformative outside gamma bursts during memory maintenance. The third assumption is that periods of activity-silence exist among selective prefrontal neural populations during memory maintenance. While the first 2 assumptions were not supported by the data, the third assumption was supported. Thus, we conclude that activity-RNN models are insufficient to explain memory maintenance in the prefrontal cortex, and alternative mechanisms, such as stp-RNNs, may be necessary to explain memory maintenance.

The absence of informative activity in the LPFC during the delay period has been observed across multiple organisms, using different methods to measure brain activity. Here, we show that the lack of elevated activity during the delay period is consistent with the sustained encoding of working memory information across selective neurons since selective neurons show increased activity to their preferred stimulus and decreased activity to their least preferred stimuli. When averaged across trials, these increases and decreases cancel each other out (Figure 2). These results are broadly consistent with Thrower and colleagues (2023) findings, who reported moderate increases in average activity during the delay period when averaging the activity of all neurons and increased decoding of working memory information in neurons with persistent activity.

We replicated the key findings reported by Lundqvist and colleagues (2016), including that informative spiking is exclusively present in electrodes that contain gamma bursts, that gamma power and gamma burst rate increased during stimulus presentation and remained elevated during the delay period, and that during gamma-burst periods the activity and information of neurons was higher than during off-periods (Figure 4). However, we also found that off-gamma periods show significant memory information (Figure 4E-F). Thus, this line of evidence does not support the existence of activity-silent periods nor stp-RNN mechanisms of working memory.

We agree with Lunkqvist colleagues (2018) that most LPFC neurons do not show sustained activity in single trials and that there are coordinated changes in activity across neurons, which may lead to high-frequency local field potential oscillations. However, activity-RNNs do not rely on single neurons with sustained activity to maintain information, nor on stable levels of activity and information across neurons (Compte et al., 2000). Instead, they rely on persistent activity across the population of selective neurons, which is consistent with our results.

It has been argued that delayed-movement tasks, such as the one performed by monkeys P and J, should not be used to study the mechanisms of working memory maintenance since they involve maintaining motor preparation information (Lundqvist et al., 2018). To address this concern, we replicated our results in a third monkey (monkey W), trained to perform a task that did not have the motor preparation confound since, during the first delay period, the monkey did not know which movement he would perform (there was a 50% chance that it would be towards the cued location and a 50% chance that it would be towards one of the other 3 locations). Our results are consistent across all 3 monkeys (suppl. Fig 1). Thus, our conclusions apply to the maintenance of information about both motor preparation and spatial working memory. In addition, if the same neurons of the LPFC encode both types of information (Tang et al., 2020), then it is reasonable to assume that the mechanism of information maintenance for motor preparation should also apply to working memory.

Due to limitations on the number of neurons we could record simultaneously, we could only explore the dependency of max-ISI on population-FR up to ∼400 Hz (Figure 4). Null-max-ISIs decreased exponentially with the population-FR. Thus, we could extrapolate what would happen with max-ISIs at much higher population-FRs if the spikes were randomly distributed. If the population-FR reaches 10,000 to 100,000 Hz, the expected max-ISI would be around 1 ms. Assuming a single cell mean firing rate of ∼20 Hz (for selective locations only), these population-FRs correspond to ∼500 to 5,000 selective neurons. The macaque LPFC has a density of ∼50,000 neurons per mm^2^ and an area of ∼20 mm^2^ (Dombrowski et al., 2001). Thus, a loose estimate of the number of neurons in area LPFC (area 9/46) is around 1 million. Given that roughly half of all neurons in the LPFC are selective during the memory delay period, for a task with 4-8 possible memory items (assuming that selectivities do not overlap), we would expect to have more than 50,000 neurons with overlapping selectivity. Thus, across the whole LPFC, the population-ISI should be lower than 1 ms if the spikes were randomly distributed.

Panichello and colleagues (2023) reported prolonged periods of chance-level decodability of working memory information in single trials. They recorded between 219 and 325 neurons simultaneously per session using high-density probes and reported that 49% of these neurons were selective during the delay period, so they recorded at most ∼170 selective neurons. If we assume that the memory fields of these neurons encompassed ∼ ¼ of the screen, they had ∼45 neurons with overlapping memory fields for every memory target. Assuming a single cell mean firing rate of ∼20 Hz (for selective locations only), their highest population-FR would be ∼900 Hz. Given this population-FR, our exponential decay model (for the null-max-ISI) would predict that the max-ISI during that session should be ∼20-30 ms. If this prediction is correct, their observation that decodability drops to chance levels for periods larger than 100 ms during the delay period of specific trials should be taken as strong evidence in favor of activity-silent mechanisms.

Here, we studied neurons recorded from FEF and LPFC separately. Even though we recorded more than twice the number of selective neurons in FEF than in LPFC (Table 1), the periods of activity-silence were longer in FEF compared to LPFC (Mann–Whitney U Test; LPFC < FEF, *p* < 0.01), suggesting that activity-silent mechanisms may be more relevant in FEF. The recurrent network of neurons that maintains a memory may encompass more than 1 brain region (e.g., FEF and LPFC combined). When we ran the same analyses on neurons simultaneously-recorded in both regions combined, we observed a similar number of activity-silent periods (Figure S2). This observation suggests that these activity-silent periods are coordinated across brain regions, which may imply a coordinating mechanism, such as top-down modulations (Miller et al., 2018).

In summary, we have found that analyzing population-ISIs provides support for the existence of activity-silent mechanisms of working memory maintenance. Our analyses showed that the long periods of chance decoding reported by Panichello and colleagues (2023) are not likely to be found by chance since, given the number of neurons that they recorded from, we would expect at most population-ISI of ∼20-30 ms. Thus, we conclude that activity-silent mechanisms are necessary to maintain working memory and/or motor preparation. However, they are seldom reflected in the spiking patterns (roughly one second of activity-silent memory maintenance every 30 minutes of memory maintenance).

## Supplementary Information

**Table S1.** The number of total units, selective units across all channels, total units recorded from gamma-modulated channels, and selective units recorded from gamma-modulated channels in each region and each monkey.

| Area | Subject | Total Units | Selective Units in All Channels | Total Units in Gmod Channels | Selective Units in Gmod Channels |
| --- | --- | --- | --- | --- | --- |
| LFPC | J | 45 | 3 | 3 | 3 |
|  | P | 75 | 34 | 34 | 34 |
|  | W | 70 | 13 | 23 | 10 |
| FEF | J | 100 | 31 | 32 | 18 |
|  | P | 12 | 4 | 9 | 4 |
|  | W | 126 | 79 | 86 | 58 |

**Figure S1.**
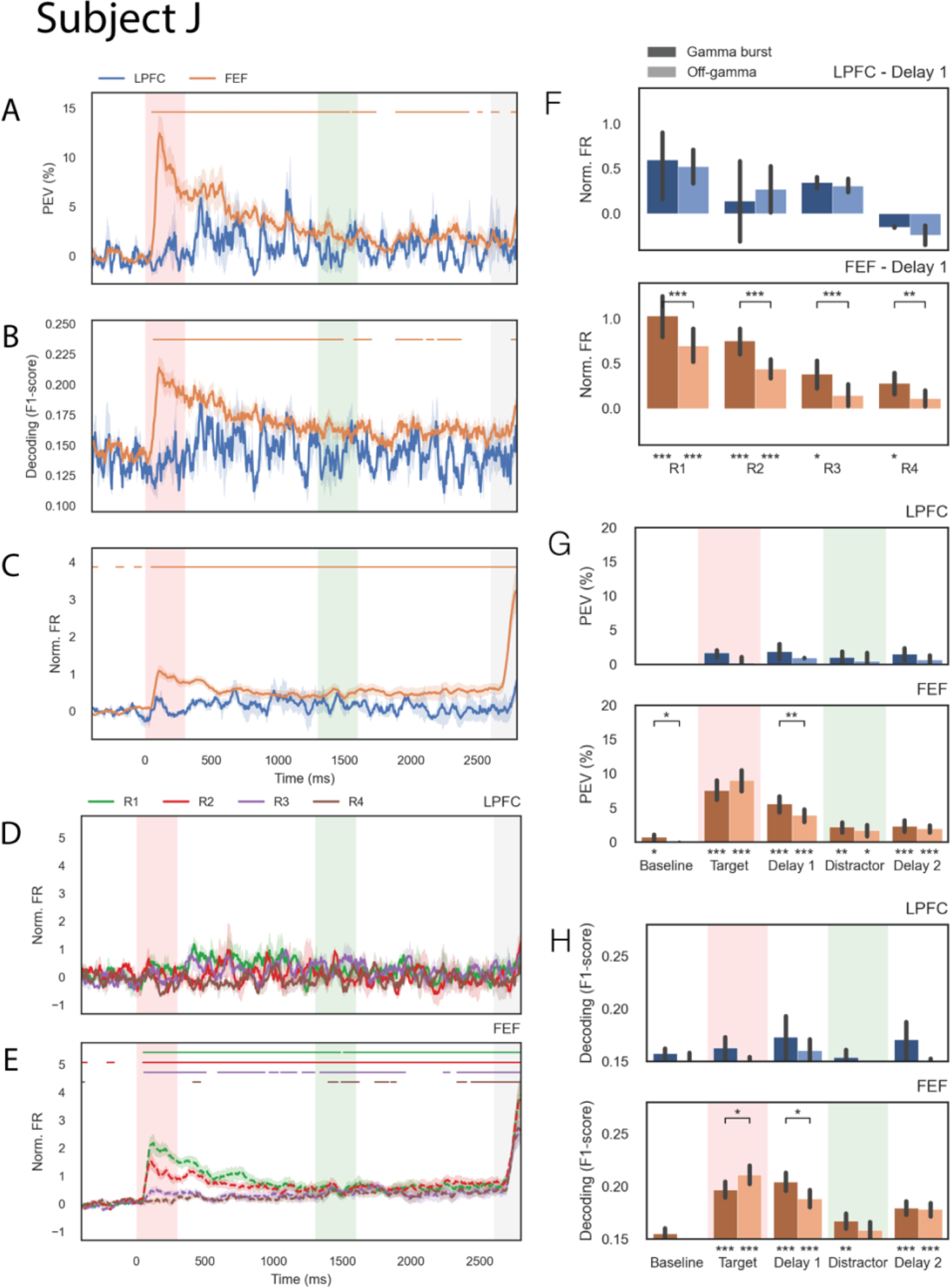
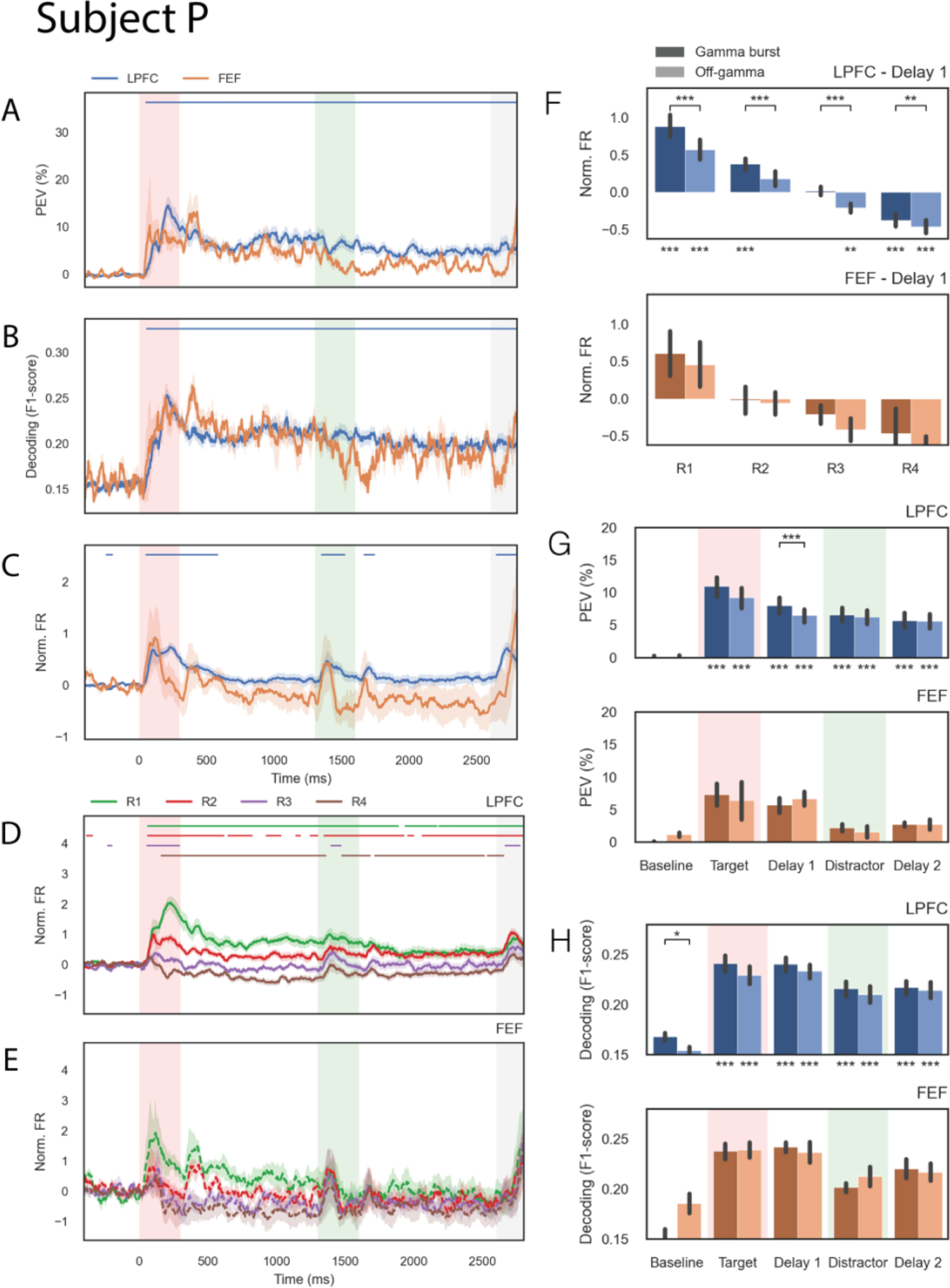
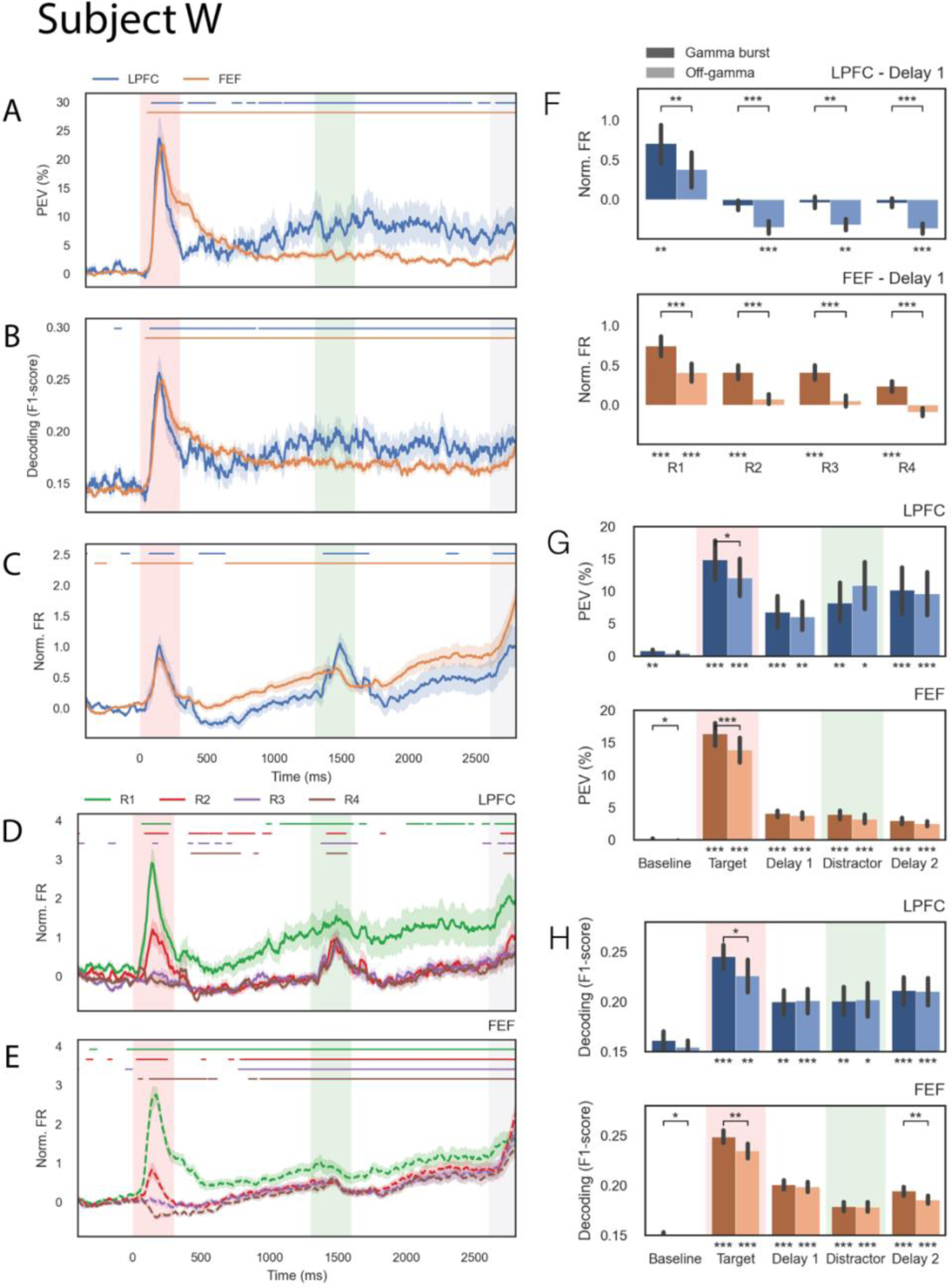
Replication of the analyses shown in Figures 2 and 3, but separated for different monkeys. Labeling and conventions are the same as for Figures 2 and 3.

**Figure S2.**
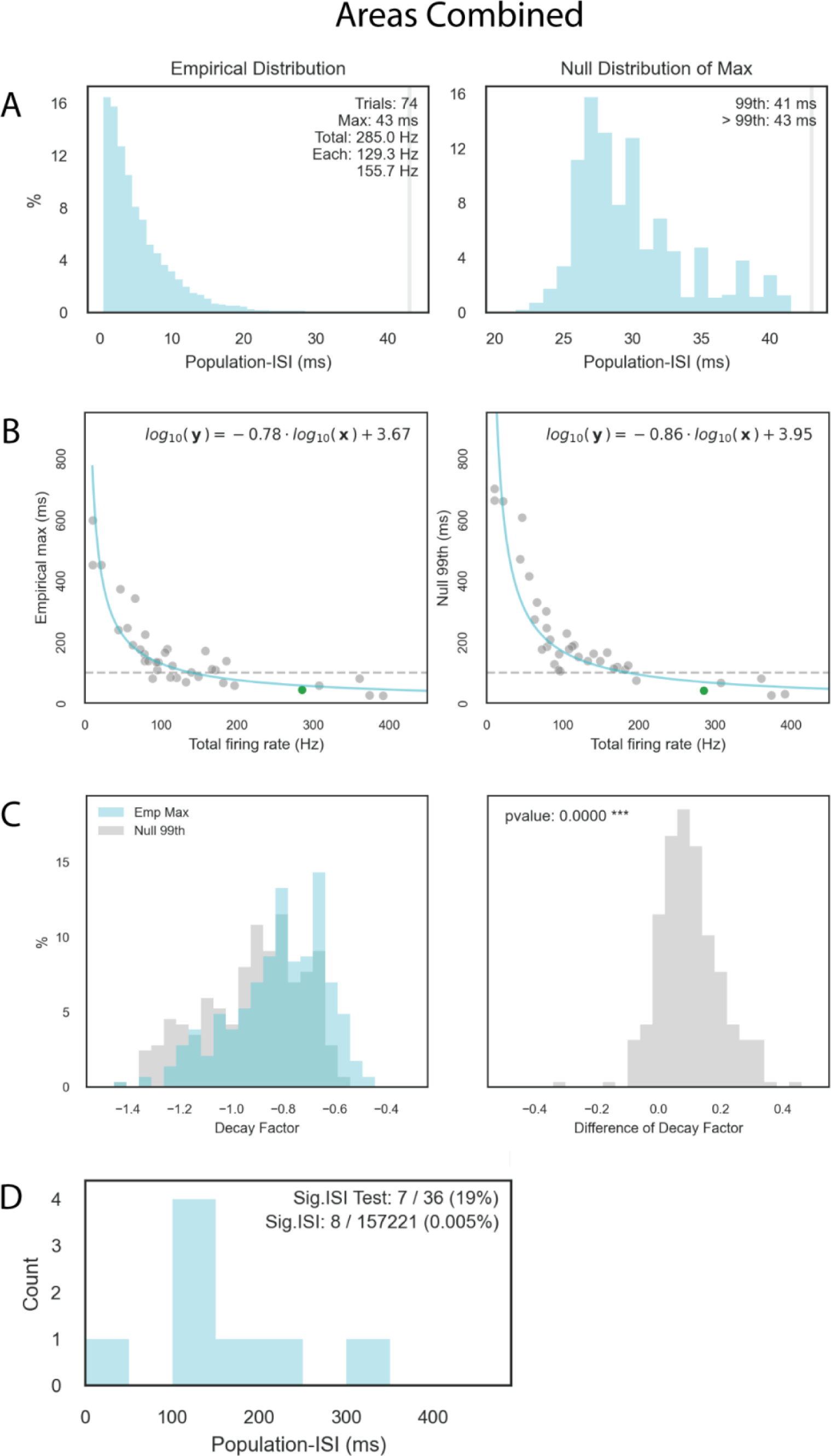
Replication of the analyses shown in Figures 4 and 5, but separated for both regions (FEF and LPFC) combined. Labeling and conventions are the same as for Figures 4 and 5.

## Notes

### Competing Interest Statement

The authors have declared no competing interest.

